# A release from developmental bias accelerates morphological diversification in butterfly eyespots

**DOI:** 10.1101/2020.04.27.063149

**Authors:** Oskar Brattström, Kwaku Aduse-Poku, Erik van Bergen, Vernon French, Paul M. Brakefield

## Abstract

Development can bias the independent evolution of traits sharing ontogenetic pathways, making certain evolutionary changes less likely. The eyespots commonly found on butterfly wings each have concentric rings of differing colors and these serially repeated pattern elements have been a focus for evo-devo research. In the butterfly family Nymphalidae, eyespots have been shown to function in startling or deflecting predators, and to be involved in sexual selection. Previous work on a model species of Mycalesina butterfly, *Bicyclus anynana*, has provided insights into the developmental control of the size and color composition of individual eyespots. Experimental evolution has also shown that the relative size of a pair of eyespots on the same wing surface is highly flexible, whereas they are resistant to diverging in color-composition, presumably due to the underlying shared developmental process. This fixed color-composition has been considered as a prime example of developmental bias with significant consequences for wing pattern evolution. Here we test this proposal by surveying eyespots across the whole subtribe of Mycalesina butterflies and demonstrate that developmental bias shapes evolutionary diversification, except in the genus *Heteropsis* which has gained independent control of eyespot color-composition. Experimental manipulations of pupal wings reveal that the bias has been released through a novel regional response of the wing tissue to a conserved patterning signal. Our study demonstrates that development can bias the evolutionary independence of traits, but it also shows how bias can be released through developmental innovations, thus allowing rapid morphological change, facilitating evolutionary diversification.

## Main text

The developmental mechanisms that generate morphology can in theory bias the independent evolution of traits sharing ontogenetic pathways, making certain evolutionary changes less likely than others (1–6). Eyespots are concentric circular markings, often with contrasting colors, that are found on the wings of many Lepidoptera (7–9). In the butterfly family Nymphalidae a series of similar eyespots is usually displayed towards the margin of the wings. These have been shown to function in startling or deflecting predators (10–13), and to be involved in sexual selection (14, 15). Eyespots have played an important role in the growing research field of evo-devo both because of their simple two-dimensional structure and their rich interspecific diversity in size and color-composition (7, 16–19). Surgical damage (20) and grafting (16, 21) experiments on early pupae demonstrated the role of the central focus in producing a signal that induces the surrounding cells to form the differently pigmented scales of an eyespot. Other studies have provided insights into the developmental control of the size and color-composition of individual eyespots (8, 18, 22). Experimental evolution using a model species of Mycalesina butterfly, *Bicyclus anynana*, has also shown that the relative size of eyespots on the same wing surface is highly flexible with little or no bias (23), whereas they are resistant to diverging in color-composition, presumably due to the underlying shared developmental process (24).

Together these studies contributed to a model of eyespot formation that, in its simplest form, involves the early pupal focal cells releasing a signal (e.g. a diffusible morphogen) that spreads out to form a circular concentration gradient (19, 24). The surrounding cells have sensitivity thresholds that direct the color of the pigmented scales that are formed. Below a certain signal threshold, the cells do not respond and will continue to develop into the normal base color of the wing, effectively forming the outer edge of the eyespot. Despite two decades of research gradually unravelling the genetic basis of the developmental processes of butterfly eyespots, the exact nature of the focal signal remains unknown. In Old World tropical butterflies of the Nymphalidae subtribe Mycalesina (25) (containing the genus *Bicyclus* and nine other genera), the order of colors typically displayed from the center (high signal level) to the outer (low signal level) ring of a normal eyespot are white, black and yellow-gold-orange (subsequently called ‘yellow’) (Fig. 1). By changing the signal level produced by each focus, the size of the individual eyespots can be modified, however it seems that the signal thresholds for the different color transitions may be fixed across the whole surface of each wing, explaining why relative proportions cannot be modulated at the level of individual eyespots (24). This has been considered as a prime example of developmental bias with significant consequences for wing pattern evolution (24).

A study investigating eyespots across a large number of Mycalesina species found that, with regard to their total size, there seems to be no strong bias limiting the independence of the main dorsal forewing eyespots (26), just as indicated by the selection experiment in *B. anynana* (22). However, to date, the question of the extent of bias in eyespot color-composition across Mycalesina has not been addressed.

**Fig. 1.**
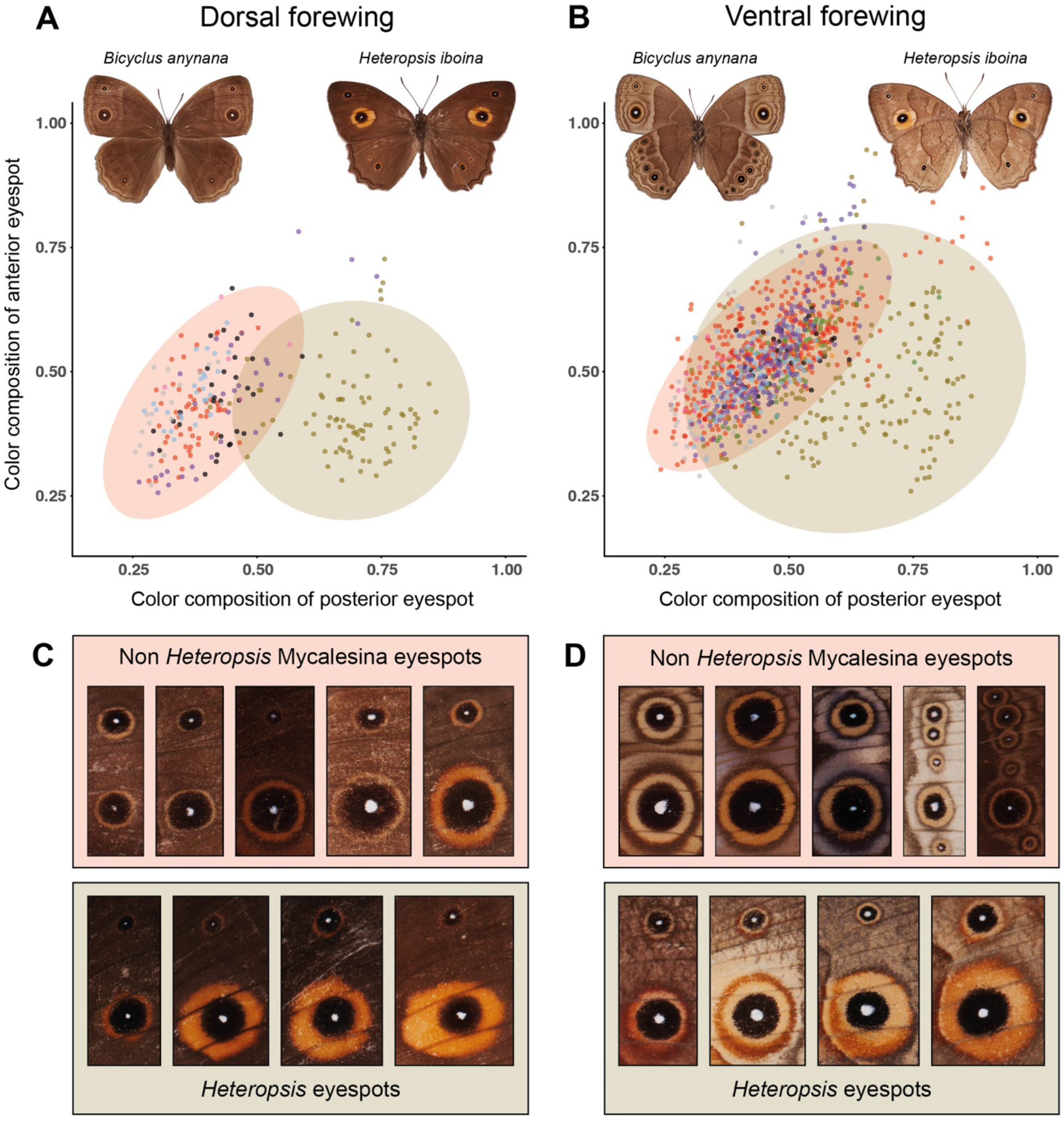
Color-compositions of dorsal and ventral forewing eyespots. Correlation of the colorcomposition – the proportion of the eyespot covered by yellow scales – between the anterior (cell M_1_) and posterior (cell CuA_1_) eyespots on the forewings of Mycalesina butterflies, as measured from 1249 male specimens from 288 taxa. The samples from different genera are color-coded following the color scheme from Fig. 2, with each dot representing an individual specimen. Computed normal confidence ellipses highlight the wide exploration of morphospace for the Malagasy genus *Heteropsis* (light olive). In contrast, the eyespot color-compositions in the other genera (light red) are strongly correlated, supporting the presence of developmental bias. Data for both (*A*) dorsal and (*B*) ventral wing surfaces only include specimens that possess both of the investigated eyespots, with both having yellow rings. (*C, D*) show examples of eyespots from species in the two color-coded groups shown in (*A*) and (*B*). (*C*) Dorsal eyespots in top panel (left to right): *Bicyclus jacksoni, Telinga misenus, Mydosama asophis, Mycalesis madjicosa* and *Brakefieldia perspicua*. Dorsal eyespots in lower panel: *Heteropsis angulifascia, H. pauper, H. ankova* and *H. turbans*. (*D*) Ventral eyespots in top panel: *Lohora dexamenus, Mydosama duponchelii, Bicyclus rileyi, Hallelesis halyma* and *Culapa kina*. Ventral eyespots in lower panel: *Heteropsis ankaratra, H. fraterna, H. strigula* and *H. turbata*.

## Results and discussion

Based on a multi-gene phylogeny containing over 80% of the described species of Mycalesina, we measured the size of separate color elements of the two main forewing eyespots from multiple male specimens from taxa for which we also had phylogenetic information. We deliberately focused on the male patterns as females of some species can be difficult to identify. The patterns in the relative size of the forewing eyespots showed no evidence of any strong bias limiting the independence of the main dorsal forewing eyespots (*SI Appendix*, Fig. S1) which is consistent with previous studies in *B. anynana* (22, 26). We then calculated the color-composition, defined as the relative proportion of yellow scales, in the two main eyespots on both dorsal and ventral forewing surfaces. There was a tight correlation between the color-composition of the two eyespots in most sampled specimens, consistent with a developmental bias limiting the independent evolution of eyespot colorcomposition. However, the majority of species in the Malagasy genus *Heteropsis* show greatly increased yellow rings in the posterior, but not the anterior, eyespot (Fig. 1), suggesting a release from the bias within this lineage.

To be able to study the correlation of eyespot color-composition in detail we calculated an index of eyespot similarity (see Methods for details) between the two main eyespots on each surface of the forewing. Using variable-rates models in the software Bayes Traits V3.0.1 (27) we estimated the ancestral states of the eyespot similarity and reconstructed historical shifts in the speed of morphological changes across the phylogeny. These analyses showed a marked increase in morphological evolution early in the genus *Heteropsis*, with only the first basal clade behaving like most other Mycalesina in showing very little evidence for significantly increased rates of evolution (Fig. 2 and *SI Appendix*, Fig. S2). The topology of the tree itself shows that there has been a release from the bias, and not just a shift to a new fixed proportion between eyespots, as the latter would have resulted in a tree with a single long branch leading up to a renewed slowly evolving phase of relative stasis (28). Once the bias is removed, active natural selection for specific morphologies, as well as neutral drift, is expected to keep the ratio between the two eyespots in a state of dynamic change as long as strong stabilizing selection is not acting against it.

**Fig. 2.**
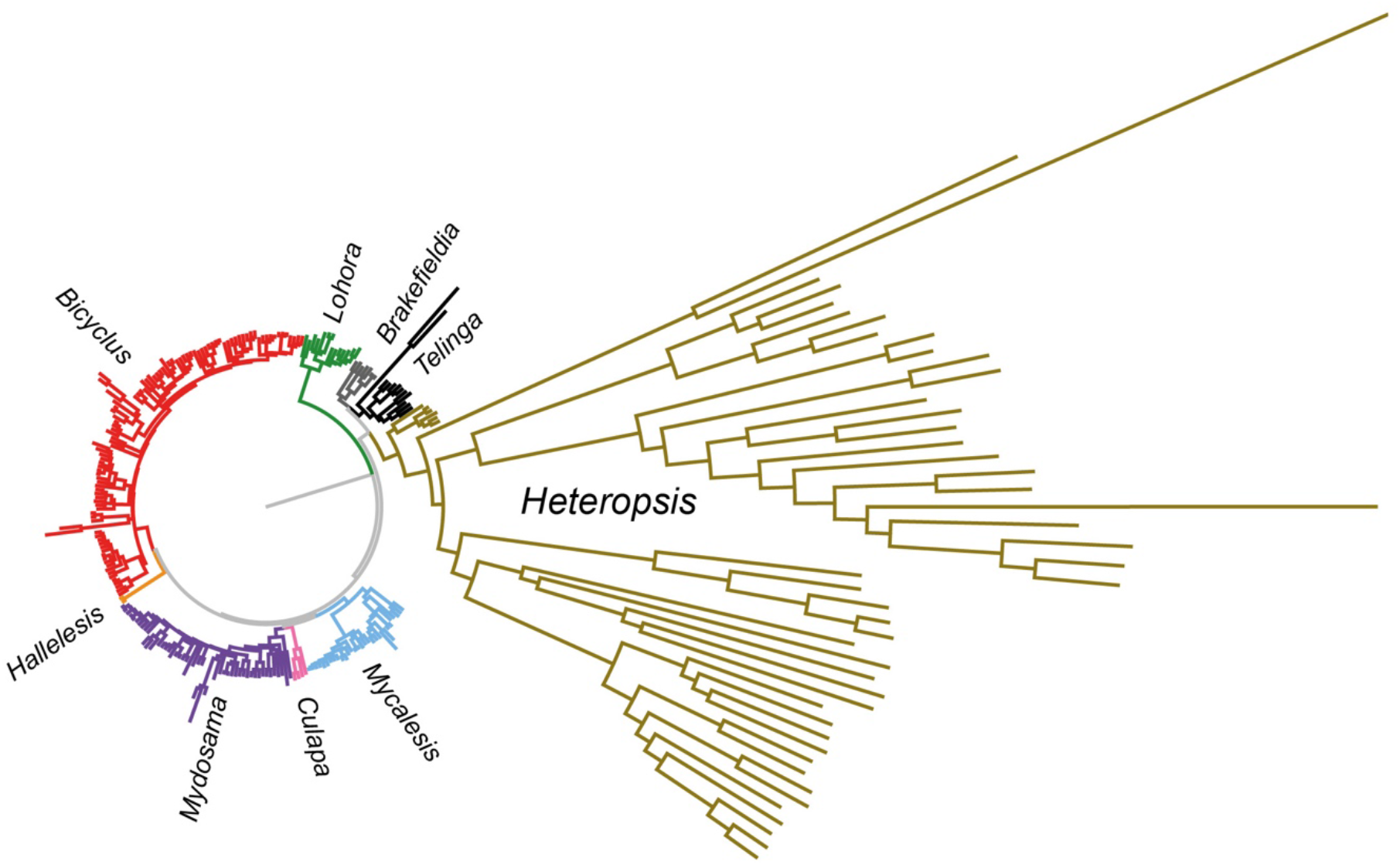
Variable rates of evolution in eyespot similarity across the Mycalesina phylogeny. Estimated historical rate of morphological evolution in eyespot similarity – the difference in color-composition of the two main eyespots – on the ventral wing surface. All branches of the Mycalesina phylogeny are scaled by the amount of morphological change estimated by the variable-rates model (Bayes Traits V3.0.1). Branches with high rates of morphological change are extended, while those with low rates are compressed. Continued acceleration of morphological diversification is observed in the genus *Heteropsis*, starting around the time of the first basal branching event, resulting in a wider exploration of morphospace (Fig. 1). The reconstruction of eyespot similarity on the dorsal forewing surface shows a similar pattern (*SI Appendix*, Fig. S2).

To investigate the developmental basis of the enlarged yellow elements of the posterior eyespots of *Heteropsis* we performed surgical grafts on developing pupae of the species *H. iboina*. These involved moving the focal cells from the large posterior eyespot into five more anterior host positions (Fig. 3*A*). This was done to investigate whether the enlarged yellow ring of the posterior eyespot resulted from the nature of the posterior focal signal, or from different response thresholds in the anterior and posterior parts of the wing. The results from 473 grafted pupae reveal that the posterior focal signal forms an ectopic eyespot with a narrow yellow ring when positioned anteriorly, but with a broader yellow region when it extends more posteriorly on the wing (Fig. 3*B*–*G*). This demonstrates that the evolutionary change that broke the linkage between eyespots has happened by modulating the response of the posterior tissue to an unchanged focal signal. Detailed inspection of the resulting wings suggested that there is a sharp border on the wing around the region of vein M_3_ with the yellow scales reaching much further down towards the posterior part of the wing once the focal signal had crossed the vein (Fig. 3*D*, *E*). Smaller ectopic eyespots, where the focal signal did not cross vein M_3_, looked like the normal anterior eyespot (in wing cell M_1_), with a narrow outer yellow ring (Fig. 3*F*). The effect was analyzed by calculating the ratio of the posterior width of the yellow ring divided by its distal width: this ratio was significantly higher for eyespots that crossed over vein M_3_ (Fig. 3*H*).

**Fig. 3.**
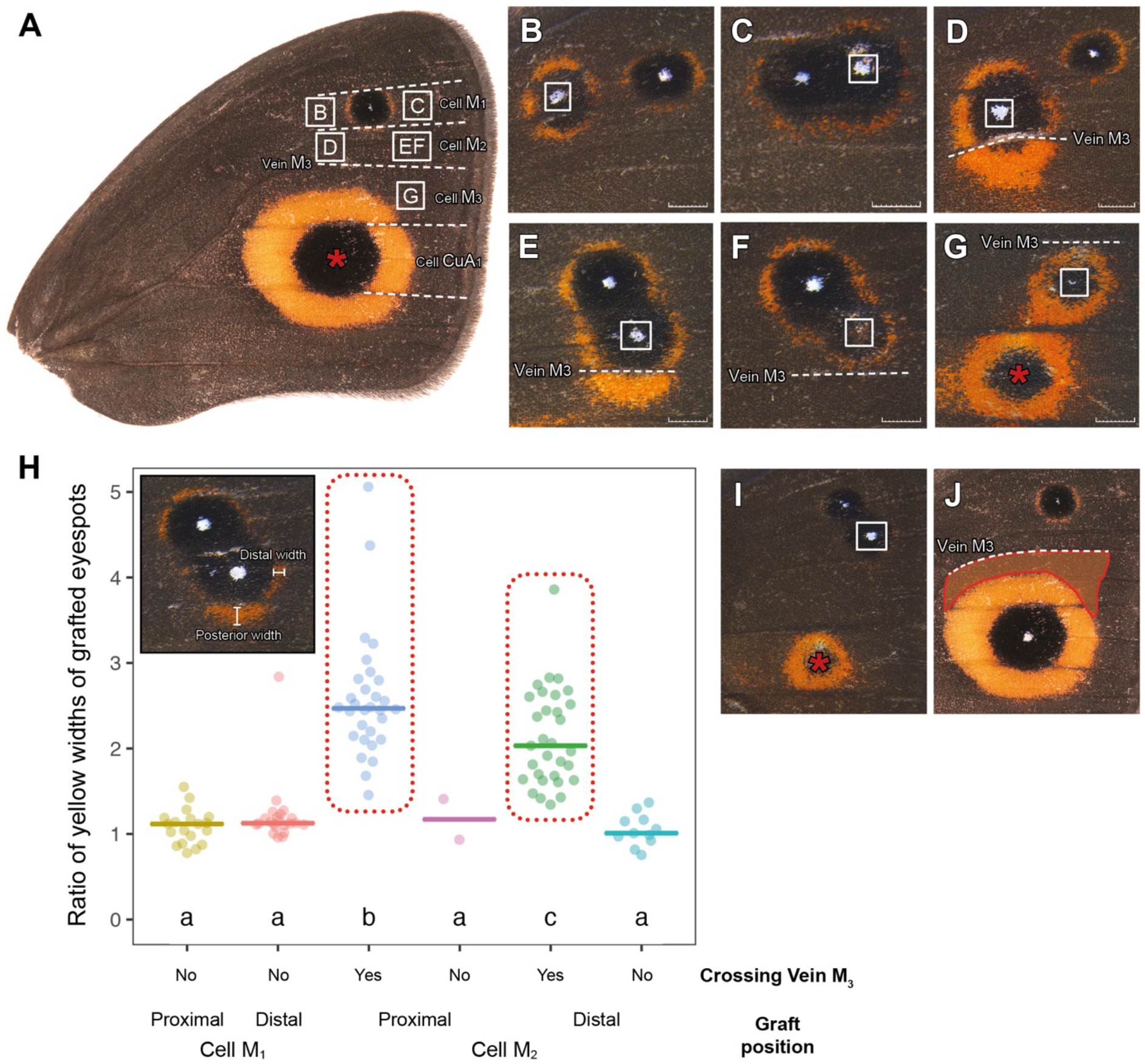
Results of grafting experiments. (*A*) The developmental basis of the enlarged yellow ring of *Heteropsis* was studied by performing surgical manipulations on pupal wings from the Malagasy butterfly *H. iboina* (normal dorsal forewing depicted). Ectopic eyespots were induced by grafting the dorsal posterior focal cells from wing cell CuA_1_ (red asterisk) into three distal and two proximal donor positions on the anterior wing (white squares). Grafts moved into a proximal (*B*) or distal (*C*) position in wing cell M_1_ – next to the normal anterior focus – induce an ectopic eyespot with a narrow yellow ring. Grafts moved into a proximal (*D*) or distal (*E, F*) position in wing cell M_2_ induce an ectopic eyespot with a narrow yellow ring that expands posteriorly when the focal signal crosses vein M_3_ (*D, E*). Grafts moved into a distal (*G*) position in wing cell M_3_ induce an ectopic eyespot with an enlarged yellow ring that typically fuses with a (reduced) posterior eyespot formed in wing cell CuA_1_ (see main text). Scale bars correspond to 1 mm. (*H*) Comparison of the ratios of the posterior and distal width of yellow in ectopic eyespots around the grafted foci showed a significant difference between eyespots crossing over vein M_3_, compared to those which did not (significant differences between groups (see Methods) are indicated by different letters). (*I*) Grafting frequently resulted in a marked reduction of patterning from the area around where the graft was taken from. (*J*) Ectopic eyespots form enlarged yellow rings within a region that is well outside the yellow area of an un-manipulated posterior eyespot (shown by overlay).

On the experimental wings, the ectopic eyespot induced around the grafted focus was usually considerably smaller than the normal posterior eyespot, and a prominent yellow (or black and yellow) pattern was frequently formed in that CuA_1_ wing cell, despite the removal of the focus when grafting the wing (Fig. 3*G*). It is likely that these effects arise from focal signalling starting at (or even before) the time of pupation (16, 21), and therefore before the time of grafting. However, it was not possible to perform surgical operations on very early pupae as the cuticle needs to somewhat harden to avoid the pupae collapsing during surgery. The residual CuA_1_ pattern was frequently strongly reduced or almost absent (Fig. 3*I*), showing that the broad yellow ring is indeed the outer element of the posterior eyespot as patterned by the focal signal (rather than being a separate, novel pattern element). Further evidence comes from the fact that ectopic patterns induced by grafts placed in cell M_2_ (and crossing over into cell M_3_), as well as grafts placed in cell M_3_ consistently produce extensive yellow scales in an upper region of cell M_3_ that is typically uniformly brown (Fig. 3*J*). In other words, if the posterior pattern were formed by a typical Mycalesina eyespot superimposed on a large yellow patch, ectopic eyespots around a grafted focus should only form narrow yellow rings in all other regions of the wing.

Our results from tissue transplantations suggest various ways in which the novel eyespot morphology in the *Heteropsis* lineage may have arisen at the genetic level. Eyespot patterning is not fully understood, but there is evidence of the involvement of the Wingless, Hedgehog and Notch signaling pathways and of transcription factors including Distal-less, Engrailed, Spalt and Antennapedia (29–31). Also, butterfly larval imaginal discs, like those of Drosophila, express certain genes (e.g. *engrailed* and *Apterous*) only in specific wing regions and these could act as selector genes, modulating patterning within the eyespots developing in different locations (19). It has been suggested that the uniformity in eyespot color-composition typical of species of Mycalesina results from this feature being specified after the phase of region-specific gene expression (19). The novelty in the *Heteropsis* wing pattern that is reflected in the differently proportioned eyespots developed by the anterior and posterior wing cells in response to focal signaling may have resulted from changes in the extent or timing of the expression of selector genes in late larval imaginal discs, or from changes in the input that particular selector genes have into the processes of eyespot patterning. In either of these ways, the response to focal signaling may have become dependent on the region-specific expression of selector genes, distinguishing anterior from posterior wing regions in the level at which yellow fate changes to the background brown color, thus setting different widths of the outer ring of their eyespots. It is intriguing that the particular region around vein M_3_ (which is distant from the Anterior-Posterior compartment border) has recently been suggested to also be of developmental significance in the wings of other Lepidopteran insects (32). A fuller understanding and experimental manipulation of the region-specific selector genes will be needed to reveal the genetic basis of the novel phenotype.

Taken together, our results show that the color-composition of the eyespots of most species of Mycalesina are strongly correlated and have little flexibility in their individual evolutionary options. This suggests that co-variation through shared development can contribute to shaping diversification, but on a macro-evolutionary scale such bias can be broken and enable rapid exploration of previously inaccessible parts of morphospace, as demonstrated here by the genus *Heteropsis*. In the field, as well as in free-flying greenhouse populations, *Heteropsis* butterflies exhibit a ritualized display of rapidly opening and closing their wings on alighting, thus exposing the conspicuous dorsal eyespot to startle a potential predator. This contrasts with the primarily deflective role of eyespots in other genera (13, 33) and suggests that the novel morphology, in combination with co-option of deimatic behavior, has facilitated evolutionary diversification in the Malagasy clade. Our results emphasize that understanding how development can bias available variation in morphology can make a valuable contribution to explaining and predicting patterns of evolutionary diversification. Essentially, we now have an example where the demonstration of the potential for developmental bias in a model species has been extended to show how this is reflected in species-rich parallel radiations. In addition, we have shown that a pattern of bias, rather than a strict developmental constraint, can be released by a developmental innovation to result in a spectacular radiation in to novel phenotypic space.

## Methods

### Phylogeny construction

303 taxa representing all known genera of Mycalesina (*Bicyclus, Brakefieldia, Culapa, Devyatkinia, Hallelesis, Heteropsis, Lohora, Mycalesis, Mydosama, Telinga*) were included as the exemplar taxa for this study. Additionally, eight taxa of the Lethina subtribe were included as outgroups. Genomic DNA was extracted from abdomens or legs using Qiagen DNEasy extraction kit following the manufacturer’s protocol. A total of ten protein-coding molecular markers were amplified and sequenced: one mitochondrial (cytochrome c oxidase subunit I, *COI*) and nine nuclear (carbamoylphosphate synthetase domain protein, *CAD*; ribosomal protein S5, *RpS5*; ribosomal protein S2, *RpS2*; wingless, *wgl*; cytosolic malate dehydrogenase, *MDH*; glyceraldhyde-3-phosphate dehydrogenase, *GAPDH*; elongation factor 1 alpha, *EF-1α*; arginine kinase, *ArgKin* and isocitrate dehydrogenase, *IDH*) gene regions were amplified. DNA amplification and sequencing followed the methodology of Aduse-Poku *et al*. (34). Sequences and voucher specimens for the phylogenetic work were primarily obtained from previously published studies of Mycalesina butterflies (25, 34–40) with some new sequences procured from field work and museum collections. A complete list of all voucher specimen data and accession codes are available in *SI Appendix*, File S1. We reconstructed our phylogenies using both Maximum likelihood (ML) and Bayesian Inference (BI) methods. The ML analysis was done in IQ-TREE v.1.6.3 (41) using best partitioning scheme and best models of nucleotide substitution suggested by ModelFinder (42). The dataset was divided *a priori*, by codon positions, resulting in 30 partitions. We estimated simultaneously the BI tree and divergence times using BEAST 1.8.4 (43). The best partitioning scheme and models of substitution for the Bayesian analyses were estimated in PartitionFinder2 (44). Given the recent controversy on Lepidoptera fossils (45), we used a more recent extensive fossil-based dating framework (46) for our dating analysis. Consequently, we constrained the node corresponding to the divergence between Lethina and Mycalesina with a uniform prior encompassing the 95% credibility interval (25.1 - 44.1 Ma) estimated for this node (46). All analyses consisted of 50 million generations with a parameter and tree sampling every 5000 generations.

### Taxonomic nomenclature

References to homologous wing veins and cells follow the system proposed by R.J. Wootton (47).

### Museum specimens

An extensive data set of images was assembled by photographing specimens from ten museums and three private collections (*SI Appendix*, Table S1). For taxa for which phylogenetic data were available we aimed to include at least five specimens in measurable condition. We focused on male specimens to prevent misidentification of taxa — females of some Mycalesina butterflies cannot be identified to species level without dissection or genetic testing (48). Seasonal polyphenism is prevalent in this group of butterflies (49, 50) and to exclude the effect of developmental plasticity (i.e. high intraspecific variation in eyespot size and color-composition, *SI Appendix*, Fig. S3) we focused on specimens that showed full expression of the wet season form phenotype, avoiding aberrant males with extreme eyespot phenotypes. The excluded specimens all showed a marked increase in the amount of yellow around their eyespots (similar to the previously documented mutations in *Bicyclus anynana* such as Goldeneye (18) and since these enlarged eyespots typically merge with neighboring eyespots they could not be measured in a repeatable way. Both this aberration and the seasonal variation affected all eyespots equally, as expected from developmental bias. For nine exceedingly rare taxa, or ones not as yet fully described, we were not able to find suitable male specimens, and six additional species were excluded because homology of color pattern elements could not be inferred with certainty. The final data set included 1249 images from 288 taxa (*SI Appendix*, Table S2).

### Image acquisition

The majority of the specimens (98.3%) were photographed using a Nikon D300 SLR with an AF-S Micro NIKKOR 60mm f/2.8G ED lens set at a fixed focus distance of 7, 9 or 10 cm. Lighting was provided by using a Metz Ring Flash 15 MS-1 and all exposure settings - including flash output - were locked to pre-determined settings to ensure comparative images. Captured RAW files were developed in Adobe Photoshop CC 2018 with fixed settings. Colors and contrasts were balanced using QPcolorsoft 501 software (2.0.1.) and reference images of a QP Card 201 that were acquired using the same procedure as for specimens. The remaining 21 specimens were analyzed from a range of photographs taken with various cameras, but with a reliable scale included in the image such that size could be correctly measured.

### Image analyses

Images were analyzed using custom-made macros and the image processing package Fiji 1.0 (51) coupled to ImageJ 1.51 (52). Two different areas of each of the eyespots in cells CuA_1_ and M_1_ were quantified as freehand selections, using a pen monitor (Huion Kamvas GT-220) to enable accurate marking of fine details. The eyespots on both the dorsal and ventral surface of the forewing were included. We measured the total area of the complete eyespot including the combined areas of the central focus, black inner-disc, and the yellow outer ring. We also measured the combined area of the central-focus and the black inner disc to be able to calculate the total and relative area of the eyespots’ yellow ring. The straight-line distance between the end points of vein CuA_2_ was quantified and used as a proxy for wing size (see below). Using this approach all raw data values could be measured with high repeatability (R^2^ > 0.99) and the variables that were derived from further calculations using the raw data also showed a high repeatability (R^2^ = 0.99 - 0.81). A summary of all repeatability tests is shown in *SI Appendix*, Table S3. Raw data from the image analyses are available in *SI Appendix*, File S1.

### Data transformation for surface-specific analyses

We calculated the relative eyespot size by dividing the total eyespot area by the squared wing index (see above). This allowed us to visually inspect our data on eyespot sizes and to compare them to previously published data (26) (*SI Appendix*, Fig. S1). The color-composition of each eyespot was defined as the relative proportion covered by yellow-gold-orange scales (i.e. the outer ring), and was calculated by dividing the yellow area by the total area. The relationships between the color-composition and relative size of the eyespots were visualized using the package *ggplot2* (53) in R (54). For each individual, eyespot similarity (ES) was assessed by dividing the color-composition of the largest eyespot by that of the smallest eyespot on the same wing surface. Hence, an ES ratio of 1 reflects that the color-compositions of the two eyespots are equal, while any deviation from 1 indicates within-surface variation in eyespot color-composition. Taxa possessing only one eyespot on the forewing, which is typical for the dorsal surface in some genera, or those that had eyespots that did not show an outer yellow ring, were excluded from further surface-specific analyses. After excluding 12 taxa that had only a single or no spot with yellow scales on either wing surface, our total data set for eyespot color-composition comprised 76 and 273 taxa for the dorsal and ventral surfaces, respectively. The eyespot in cell CuA_1_ was the largest eyespot for all species included in the dorsal analyses and for all but nine of the species included in the ventral analyses.

### Evolutionary analyses

To detect shifts in the rate of eyespot similarity (ES) across the phylogeny, we used the variable-rates model described by Baker *et al*. (28) as implemented in the software package BayesTraits V3.0.1 (27) (available at www.evolution.rdg.ac.uk). All analyses were conducted using the time-calibrated, multi-gene consensus tree and mean eyespot similarity (ES) per taxon as the focal trait. Reversible-jump MCMC algorithms were run for 105 million iterations, with a burn-in period of 5 million generations, after which the chain was sampled every 100 000th iteration. Priors were kept at default for all analyses and each run was repeated 5 times to confirm the stability of the log marginal likelihoods and to check the topologies of the consensus trees. The log marginal likelihoods for the *m*_0_ (fixed-rates) and *m*_1_ (variable-rates) models were estimated using stepping-stone sampling implemented in BayesTraits, and then used to compute a log Bayes factor (BF). We ran the stepping-stone sampler with 1000 stones and 100 000 iterations for each stone following the completion of each analysis. BFs were used as the test statistic; values greater than 2 are typically considered positive evidence and BFs greater than 10 are taken as very strong evidence for rate variation (55). We found strong support for variable rates of morphological change on both wing surfaces (ventral, BF_mean_ = 160.4; dorsal, BF_mean_ = 22.6; *SI Appendix*, Table S4). The variable-rates model identified increased rates of evolution as mainly occurring along the branches of the *Heteropsis* clade (Fig. 2, *SI Appendix*, Fig. 2), strongly implying that these species have gained independent control of eyespot color-composition. Input data for the evolutionary analyses are available in *SI Appendix*, File S1.

### Grafting experiment

Larvae from a laboratory population of *Heteropsis iboina* (56) were reared on basketgrass (*Oplismenus compositus*) in climate-controlled chambers (Panasonic MLR-352H-PE) at 25°C, 75% relative humidity (RH) and a 12:12 L:D cycle. Pre-pupae were collected daily and placed in compartmentalized Petri dishes. Using time-lapse photography we recorded the time of pupation to the nearest 15 minutes. Grafting operations were performed on the dorsal surface of the forewing, 4 to 5 hours after pupation, when the pupal cuticle had hardened sufficiently to allow surgery but before the underlying wing epidermis had separated. The grafting experiment followed procedures as previously described for *Bicyclus anynana* (20). We performed five surgical manipulations, all involving moving the eyespot focus located in cell CuA_1_ (which is easy to locate on the pupal epidermis) to a new host site on the same wing. The tissue from the host site was then used to cover the focal area that was used as donor tissue. The five host sites included three distal (in relation to the location of the eyespot in cell M_1_) positions in cells M_1_, M_2_ and M_3_, as well as two proximal locations in cells M_1_ and M_2_ (see Fig. 3 in main text). After grafting, pupae were left untouched for 30 minutes to allow the hemolymph to seal the incision sites. Subsequently, pupae were placed in individual transparent pots and returned to the climate-controlled chambers to complete development. One day after eclosion the adults were frozen to −18 °C, after which the wings were removed using surgical scissors. The dorsal surface of the experimental wings was photographed using a Leica DFC495 digital camera coupled to a Leica M_1_25 stereomicroscope. The numbers of performed grafts and associated eclosion rates are presented in *SI Appendix*, Table S5. Images were inspected by eye to assess the effect of the grafting procedure. To estimate the effect of grafts crossing vein M_3_ we measured the distal and posterior width of ectopic eyespots showing clear yellow outlines in grafts placed in cell M_1_ and M_2_ (N=115). A ratio was calculated for each eyespot by dividing the posterior width by the distal width. These ratios were used as the dependent variable in a linear model with the graft position and the crossing of the vein as a concatenated fixed factor (fig 3h). Post hoc pairwise comparisons (Tukey’s HSD; α = 0.05) were performed using the *emmeans* package (57).

## Data availability

Raw data from the image analyses, input data for the evolutionary analyses, and information on all voucher specimens used in the phylogeny is available as *SI Appendix*, File S1. The phylogenetic consensus tree used for the evolutionary analyses is available as *SI Appendix*, File S2. All other data are available from the corresponding author upon request.

## Acknowledgements

Photographs of analyzed specimens were taken from the museum collections of ABRI (Nairobi), MGCL (Gainesville), MNHB (Berlin), MRAC (Tervuren), NHMUK (London), NHMW (Vienna), NHRS (Stockholm), OUMNH (Oxford), NBNC (Leiden), SMNS (Stuttgart) and the personal research collections of Oskar Brattström, Peter Roos and Robert Tropek. We thank all the Curators and the Trustees of these collections. Rob Freckleton and Michael Akam provided comments on the manuscript. Elishia Harji assisted with proofreading and language editing. Simon Chen photographed specimens from the collections at SMNS. This work was funded by grants from the John Templeton Foundation (#60501) and the European Research Council (EMARES #250325) to P.M.B.

## Author contributions

O.B., V.F., and P.M.B. conceived the study and designed the experiment. O.B. identified and photographed museum specimens, processed the images and performed the grafting experiments. K.A.P. constructed the phylogeny, collected the *H. iboina* laboratory population and performed early grafting experiments. Final statistical analyses were performed by O.B. with support from E.v.B.. O.B., V.F., and P.M.B. wrote the manuscript, with contributions from K.A.P. and E.v.B.. All authors read and approved the final version of the manuscript.

---

**Figure S1.**
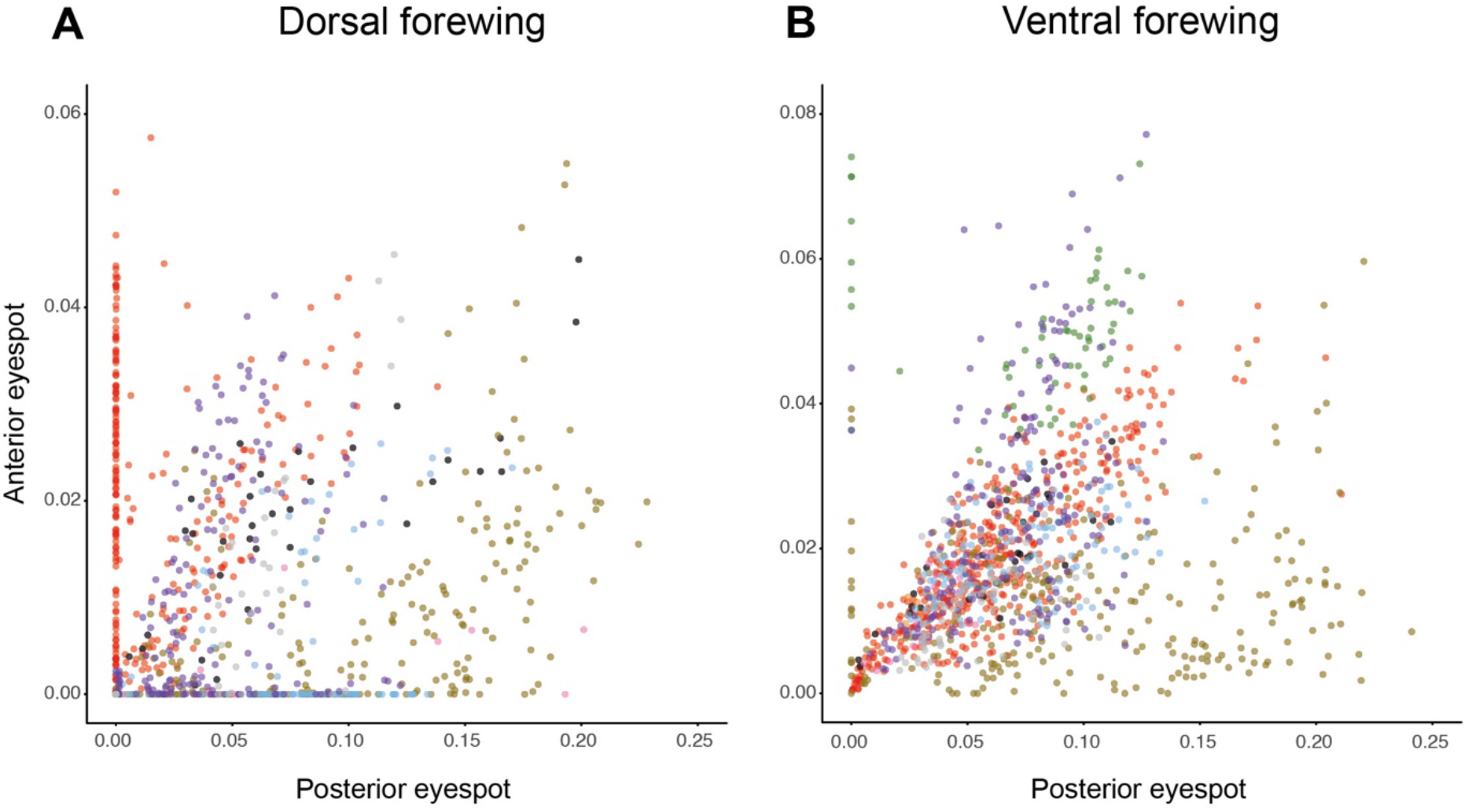
Relative size of forewing eyespots. Plots show the distribution of the total area (including all color elements from center to the outer edge of the yellow ring if present) of forewing eyespots for 1249 measured specimens of Mycalesina butterflies representing 288 taxa. The data shown in the plots is the relative area of eyespots calculated by dividing the absolute eyespot area by an index of wing size squared (see Material and Methods). (*A*) Dorsal wing surface (16 outliers not shown) and (*B*) Ventral wing surface (5 outliers not shown). The different genera are color coded following the color scheme of Figure 2 in the main text.

**Figure S2.**
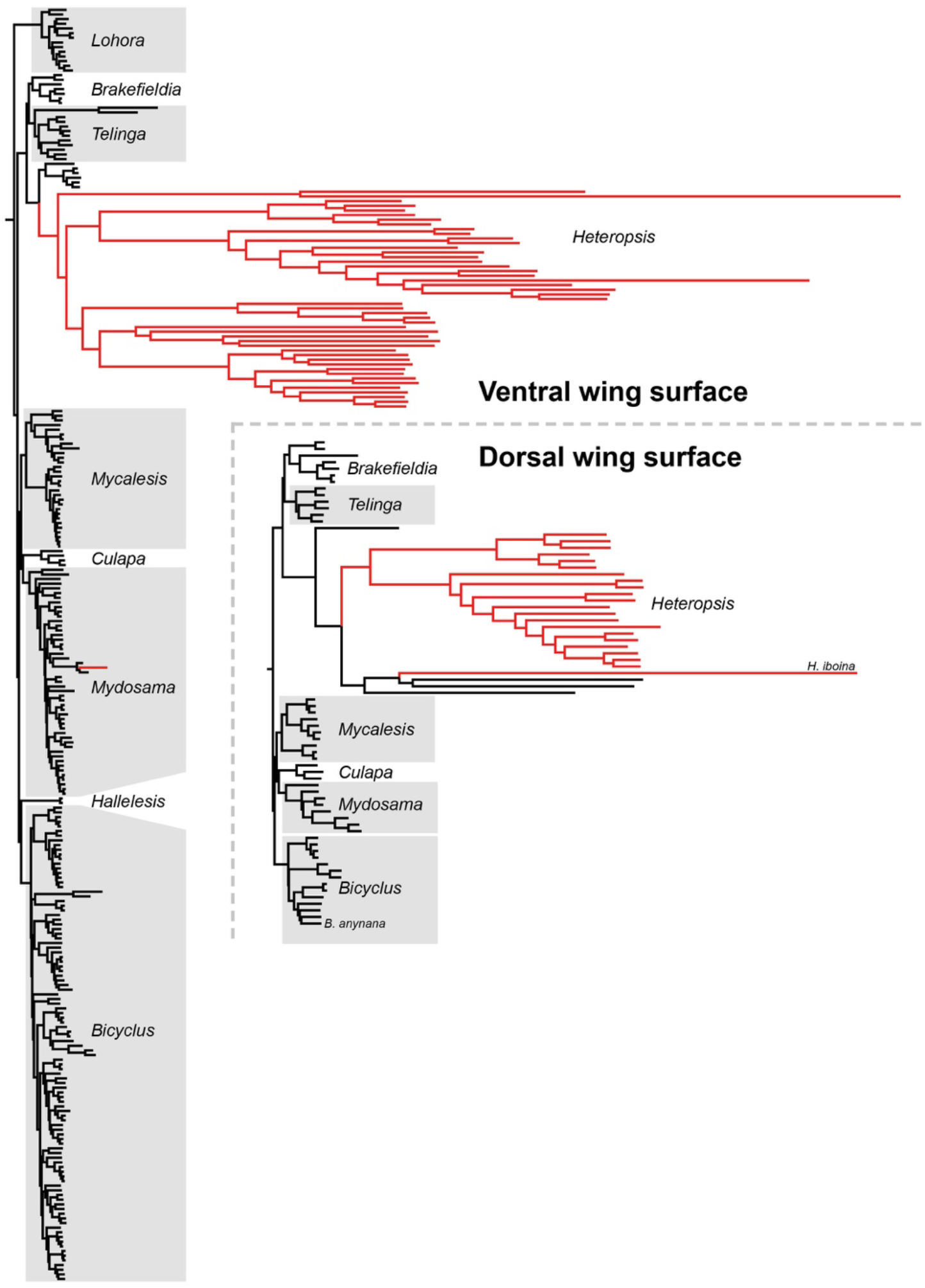
Comparison of evolutionary dynamics on dorsal and ventral wing surfaces. Estimated historical rate of morphological evolution in forewing eyespot similarity on the dorsal and ventral wing surface. Branches in red show evidence of significant positive phenotypic selection following the method suggested by Baker *et al*. (28). The data shown for the ventral surface is the same as displayed in the circular tree in Figure 2 in the main text. Sample size for the dorsal surface was reduced owing to taxa missing either one of the eyespots on this surface or because the eyespots did not contain yellow outer rings. In the dorsal eyespot phylogeny the position of the two species for which grafting experiments have been performed (*Bicyclus anynana* and *Heteropsis iboina*) are indicated.

**Figure S3.**
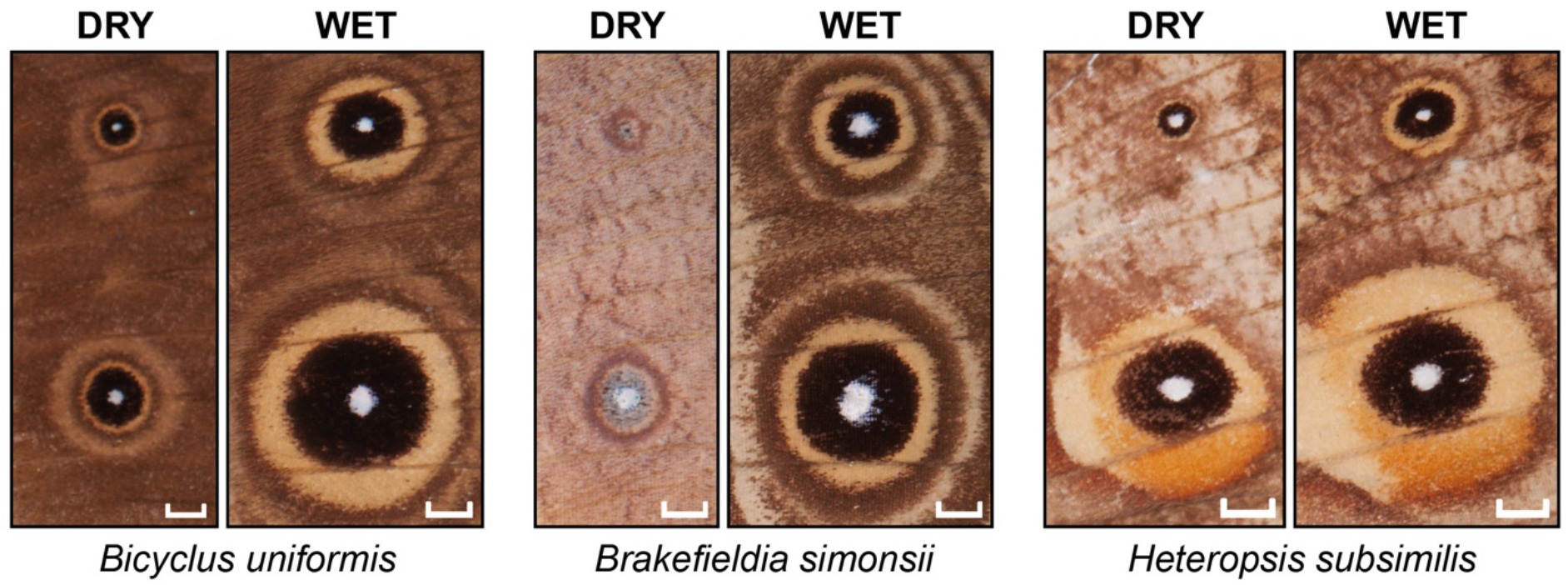
Examples of dry (left) and wet (right) season morphologies in Mycalesina eyespots. The anterior (top) and posterior (bottom) eyespots of the ventral forewing are shown from three representative species. In this study we focused on specimens that showed full expression of the wet season form phenotype. Dry season specimens have smaller eyespots, and the color-compositions can differ from those of the wet season specimens (e.g. *B. simonsii*), but there is no indication that they do not follow the same bias in color-composition in the majority of the species (e.g. *B. uniformis* and *B. simonsii*). The release from this developmental bias in the genus *Heteropsis* involves displaying a larger relative proportion of yellow in the posterior spot regardless of seasonal phenotype, and is represented by *H. subsilimis* in this figure. The small eyespots of the dry season form are harder to measure with high repeatability, not only because they represent smaller areas on the images, but also the nature of the wing scale cells makes them less reliable. Since each individual scale can take on only a single color, the smaller eyespots will have less regular outlines, and changes in single, or a few, scales can have comparatively large effects on the proportional areas. Scale bars correspond to 1 mm.

**Table S1.**
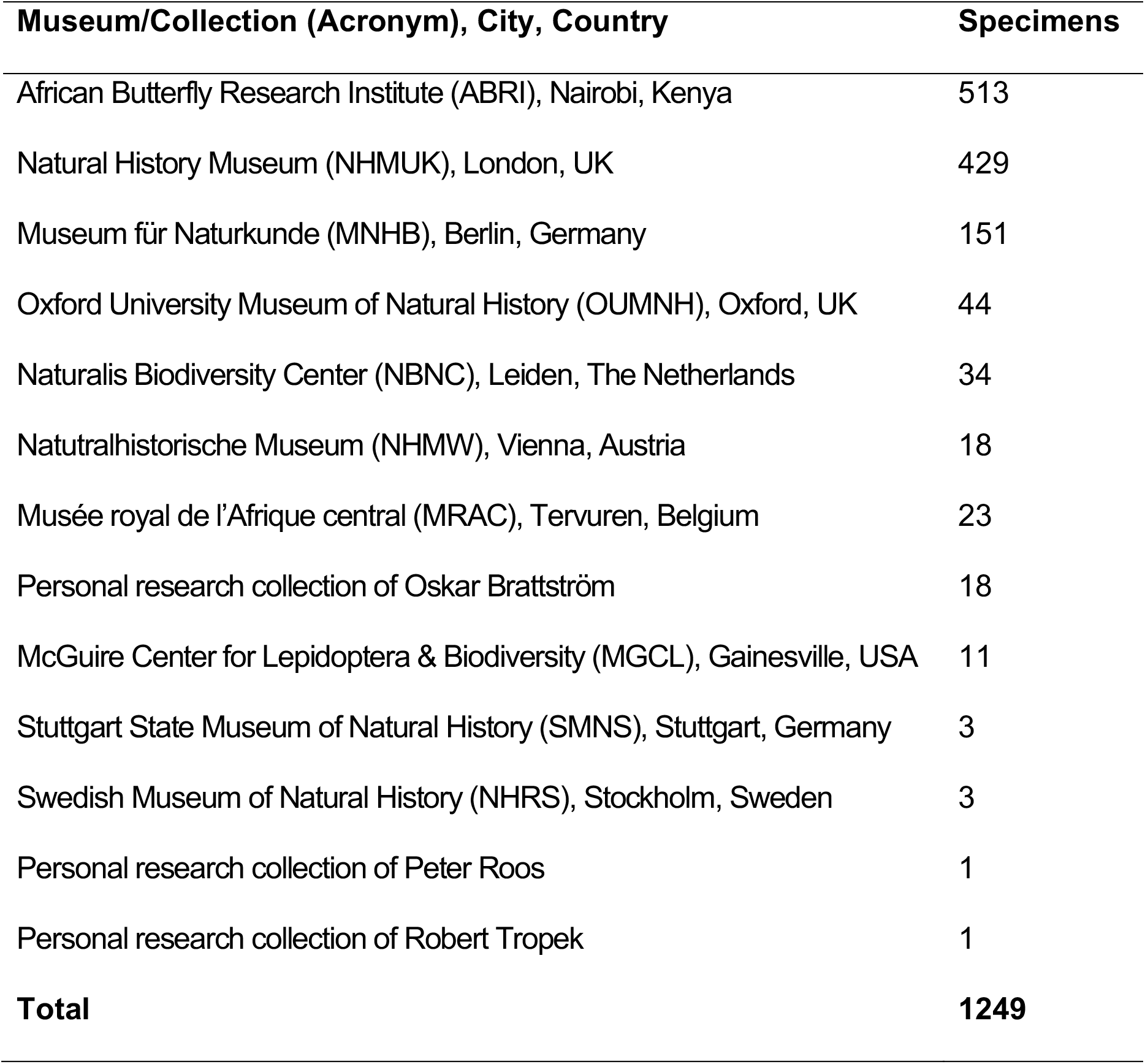
Sources of photographed specimens.

**Table S2.** Taxonomic distribution of taxa analyzed shown for all currently recognized genera of Mycalesina. Mean number of measured samples per taxa, grouped by genera, is also shown.

| **Genus** | **Total taxa** | **Species** | **Additional**<br>**subspecies** | **Undescribed**<br>**taxa** | **Total samples**<br>**(mean/taxon)** |
| --- | --- | --- | --- | --- | --- |
| *Bicyclus* | 102 | 93 | 7 | 2 | 485 (4.75) |
| *Brakefieldia* | 7 | 7 | - | - | 28 (4.71) |
| *Culapa* | 4 | 4 | - | - | 19 (4.75) |
| *Devyatkinia* | 0 | 0 | - | - | 0 (0) |
| *Hallelesis* | 2 | 2 | - | - | 10 (5.00) |
| *Heteropsis* | 64 | 62 | 2 | 2 | 241 (3.77) |
| *Lohora* | 16 | 16 | - | - | 62 (3.88) |
| *Mycalesis* | 30 | 19 | 11 | - | 127 (4.23) |
| *Mydosama* | 51 | 41 | 9 | 1 | 218 (4.27) |
| *Telinga* | 12 | 12 | - | - | 54 (4.50) |
| **Total** | **288** | **254** | **29** | **5** | **1249 (4.34)** |

**Table S3.** Summary of repeatability tests performed by remeasuring the ventral spots from 50 specimens picked at random from the full dataset. The correlation coefficient and r^2^ values for the correlations between original and remeasured values are given for all raw data traits (bold) and variables calculated from the raw data (normal text).

| **Measurement** | **coeff.** | **$r^2$** |
| --- | --- | --- |
| **Total Area M<sub>1</sub>** | **0.983** | **1.00** |
| **Black + Focus Area M<sub>1</sub>** | **0.994** | **1.00** |
| Yellow Area M<sub>1</sub> | 0.949 | 0.98 |
| Yellow Proportion M<sub>1</sub> | 1.004 | 0.93 |
| **Total Area CuA<sub>1</sub>** | **0.994** | **1.00** |
| **Black + Focus Area CuA<sub>1</sub>** | **0.967** | **1.00** |
| Yellow Area CuA<sub>1</sub> | 1.021 | 0.99 |
| Yellow Proportion CuA<sub>1</sub> | 1.011 | 0.94 |
| **Size Index** | **1.011** | **1.00** |
| Eyespot Similarity Index | 0.853 | 0.81 |

**Table S4.** Mean Log Marginal Likelihood for the complex Variable rates model, the simple Fixed rate model, and the resulting Mean Log Bayes Factor. Values were estimated using the Stepping Stone Sampler implemented in Bayes Traits 3.1. set to sample 1000 stones with 100000 iterations. We ran five separate analyses of each model and resulting minimum and maximum values are shown in brackets.

| Wing surface | Log Marg. Lh. | Log Marg. Lh. | Log Bayes Factor |
| --- | --- | --- | --- |
|  | Variable Rates | Fixed Rate |  |
| Dorsal | -12.75 [-12.84, -12.60] | -24.04 [-24.08, -24.01] | 22.6 [22.4, 23.0] |
| Ventral | 153.98 [153.84, 154.27] | 73.77 [73.71, 73.80] | 160.4 [160.2, 160.9] |

**Table S5.** Summary of grafting experiment showing number of grafts per position (one per specimen) and the number of successful eclosions with measurable effects of the transplanted tissue.

| **Donor** | **Host** | **Grafts** | **Eclosions** | **Ecl. rate** |
| --- | --- | --- | --- | --- |
| CuA<sub>1</sub> | Distal M<sub>1</sub> | 112 | 97 | 86.6% |
| CuA<sub>1</sub> | Distal M<sub>2</sub> | 112 | 95 | 84.8% |
| CuA<sub>1</sub> | Distal M<sub>3</sub> | 113 | 94 | 83.2% |
| CuA<sub>1</sub> | Proximal M<sub>1</sub> | 107 | 94 | 87.9% |
| CuA<sub>1</sub> | Proximal M<sub>2</sub> | 112 | 93 | 83.0% |
| **Total** |  | **556** | **473** | **85.1%** |

